# Individual variation in longitudinal postnatal development of the primate brain

**DOI:** 10.1101/396887

**Authors:** G. Ball, M. L. Seal

## Abstract

Quantifying individual variation in postnatal brain development can provide insight into cognitive diversity within a population and the aetiology of common neuropsychiatric and neurodevelopmental disorders that are associated with adverse conditions in early life. Non-invasive studies of the non-human primate can aid understanding of human brain development, facilitating longitudinal analysis during early postnatal development when comparative human populations are difficult to sample.

In this study, we perform analysis of a longitudinal MRI dataset of 32 macaques, each with up to five magnetic resonance imaging (MRI) scans acquired between 1 and 36 months of age. Using nonlinear mixed effects model we derive growth trajectories for whole brain, cortical and subcortical grey matter, cerebral white matter and cerebellar volume. We then test the association between individual variation in postnatal tissue volumes and birth weight.

We report nonlinear growth models for all tissue compartments, as well as significant variation in total intracranial volume between individuals. We also demonstrate that subcortical grey matter varies both in total volume and rate of change between individuals and is associated with differences in birth weight. This supports evidence that subcortical grey matter is specifically vulnerable to adverse conditions ***in utero*** and highlights the importance of longitudinal MRI analysis in developmental studies.

## Introduction

Characterising individual variation in early brain development is critical to understanding the emergence of cognitive and behavioural diversity within populations and may provide insight into the aetiology of common neuropsychiatric and neurodevelopmental disorders (Foulkes and Blakemore 2018; Seghier and Price 2018). Longitudinal data is essential to the valid estimation of individual differences (Rogosa et al. 1982; Cudeck 1996; Kanai and Rees 2011; Madhyastha et al. 2017). Longitudinal studies of brain growth using magnetic resonance imaging (MRI) in childhood have revealed several key characteristics common across samples (Giedd et al. 1999; Gilmore et al. 2012; Mills et al. 2016). Brain growth is at its most rapid in the first year of life, and accompanied by nonlinear changes to the distributions of cortical and subcortical grey matter and cerebral white matter volume that continue through childhood and adolescence (Tamnes et al. 2010, 2017; Lebel and Beaulieu 2011; Gilmore et al. 2012; Aubert-Broche et al. 2013; Holland et al. 2014; Raznahan et al. 2014; Wierenga et al. 2014). Increasing evidence suggests that neurodevelopmental variation in youth is partly mediated by the long-term impact of adverse conditions ***in utero*** and interruption to developmental processes in early life can have a long-standing impact on cerebral development and cognitive function (Morsing et al. 2011; Raznahan et al. 2012; Walhovd et al. 2012; Sucksdorff et al. 2015; Ball et al. 2017). Alterations to typical trajectories of brain development are implicated in neurodevelopmental disorders such as autism spectrum disorder, attention-deficit hyperactivity disorder and schizophrenia and may be rooted in adversity in early life (Courchesne et al. 2003; Douaud et al. 2009; Morsing et al. 2011; Lampi et al. 2012; Chiapponi et al. 2013; Hazlett et al. 2017). It is therefore vital to understand how the brain development varies over time within individuals, and how early disturbances to foundational processes may impact this trajectory.

Both human and non-human primates undergo a protracted period of postnatal cerebral development, characterised by rapidly increasing cerebral volume, expansion of higher-order neocortical areas and axonal myelination that coincide with dynamic alterations in gene expression (Kinney et al. 1988; Bourgeois et al. 1994; Amlien et al. 2016; Bakken et al. 2016). Across mammalian species, the relative timing of key neurodevelopmental events including neurogenesis, synaptogenesis and pruning, postnatal myelination, and brain growth are relatively conserved (Workman et al. 2013), and the relationship between brain size and body size are largely governed by allometric scaling laws across both ontogeny and phylogeny (Passingham 1985; Harvey and Pagel 1988). Despite this, several features unique to the primate brain development have been identified, including expanded cellular proliferation during neurogenesis (Smart et al. 2002) and increased cortical neuronal density for a given brain size (Herculano-Houzel et al. 2014). Non-invasive studies of the non-human primate brain with magnetic resonance imaging (MRI) can therefore provide useful insight into human brain development, facilitating longitudinal analysis and cross-species comparison, particularly during early postnatal development when comparative human populations are difficult to sample (Hunsaker et al. 2014; Scott et al. 2016; Grootel et al. 2017; Sakai et al. 2017).

As with humans, longitudinal MRI analyses in macaques have found that total brain volume approaches two-thirds of full adult volume by one week of age, and follows a nonlinear trajectory over time with an initial rapid growth rate that slows from around 4 months onwards (Malkova et al. 2006; Payne et al. 2010; Scott et al. 2016). In both species, total brain volume is, on average, larger in males, and postnatal growth is largest in frontal, parietal and temporal lobes with similar trajectories in subcortical and cerebellar volumes (Gilmore et al. 2012; Scott et al. 2016). In humans, longitudinal analyses of structural MRI have found that cortical grey matter volume doubles in the first year of life and peaks in early childhood, then gradually decreases during adolescence; white matter, on the other hand, follows a more stable trajectory with age (Gilmore et al. 2012; Mills et al. 2016). Using manual tissue segmentation, Malkova et al. found that white matter volume undergoes rapid increase up to 5 months, followed by a period of relatively stable linear growth until 4 years of age (Malkova et al. 2006). In macaques, longitudinal analyses of postnatal cortical grey matter volume are lacking, however in a cross-sectional study of 37 juvenile macaques, Knickmeyer et al. reported linear and quadratic increases of total brain volume and white matter volume, respectively, but no significant age-related change in cortical grey matter (Knickmeyer et al. 2010).

Despite relative concordance between sites and studies on the typical growth trajectory of primate brain development, open questions remains on how best to define brain tissue volume change over time – i.e.: is the observed trend linear, quadratic, or cubic with age? Nonlinear growth implies periods of accelerated growth that may represent critical developmental periods or windows of vulnerability to injury. In humans, several studies have reported developmental peaks in cortical grey matter volume thought to define important points of inflection in brain development that vary meaningfully between sexes and across clinical cohorts (Sowell et al. 2004; Lenroot et al. 2007; Shaw et al. 2007; Raznahan et al. 2011; Alexander-Bloch et al. 2014). Similar observations have been made for subcortical grey matter and cerebellar volume (Raznahan et al. 2014; Wierenga et al. 2014). However, the choice of functional form can alter when, and if, peaks are defined, and the best-fit model can be dependent on the sampled population, image preprocessing strategies, model selection strategy or study design (Mills and Tamnes 2014; King et al. 2017; Walhovd et al. 2017).

Generalised additive models (GAMs) offer an alternative approach to polynomial modelling for nonlinear trends (Hastie and Tibshirani 1986). In GAMs, linear predictors are replaced by smooth functions, with the degree of smoothness estimated during model fitting (Wood 2011). As such, GAMs offer an efficient and flexible non-parametric alternative to linear models that do not require the specification of a functional form (quadratic, cubic, etc.) for the relationship between tissue volume and age (Gennatas et al. 2017; Herting et al. 2018).

In this study, we use GAMs to examine trajectories of postnatal brain development in the macaque. Using data from an open-source, neurodevelopmental macaque database (Young et al. 2017), we extend upon previous reports and model nonlinear growth of cortical grey matter, white matter, subcortical grey matter and the cerebellum. We quantify both group averaged trajectories and the individual variation present within the sample and report associations between birth weight and postnatal tissue volume.

## Methods

### Data

Longitudinal MRI data were acquired from the UNC-Wisconsin Rhesus Macaque Neurodevelopment Database. (Young et al. 2017) This publicly available database contains MRI scans from 34 rhesus macaques (***macaca mulatta***) housed at the Harlow Primate Laboratory (HPL) at the University of Wisconsin-Madison and scanned up to five times between the ages of 2 weeks and 36 months. The research protocol was approved by the local Institutional Animal Care and Use Committee. Data are available to download from: https://data.kitware.com/#collection/54b582c38d777f4362aa9cb3. Full study protocols are detailed in Young et al. (2017).

After quality control and image processing (see below), 9 scans were excluded, resulting in a final dataset comprising 32 macaques (18 male) with between 2 and 5 scans (mean number of scans = 4.7) and scanned between 1 and 36 months (mean age at scan=13.3 months)(Figure 1).

**Figure 1:**
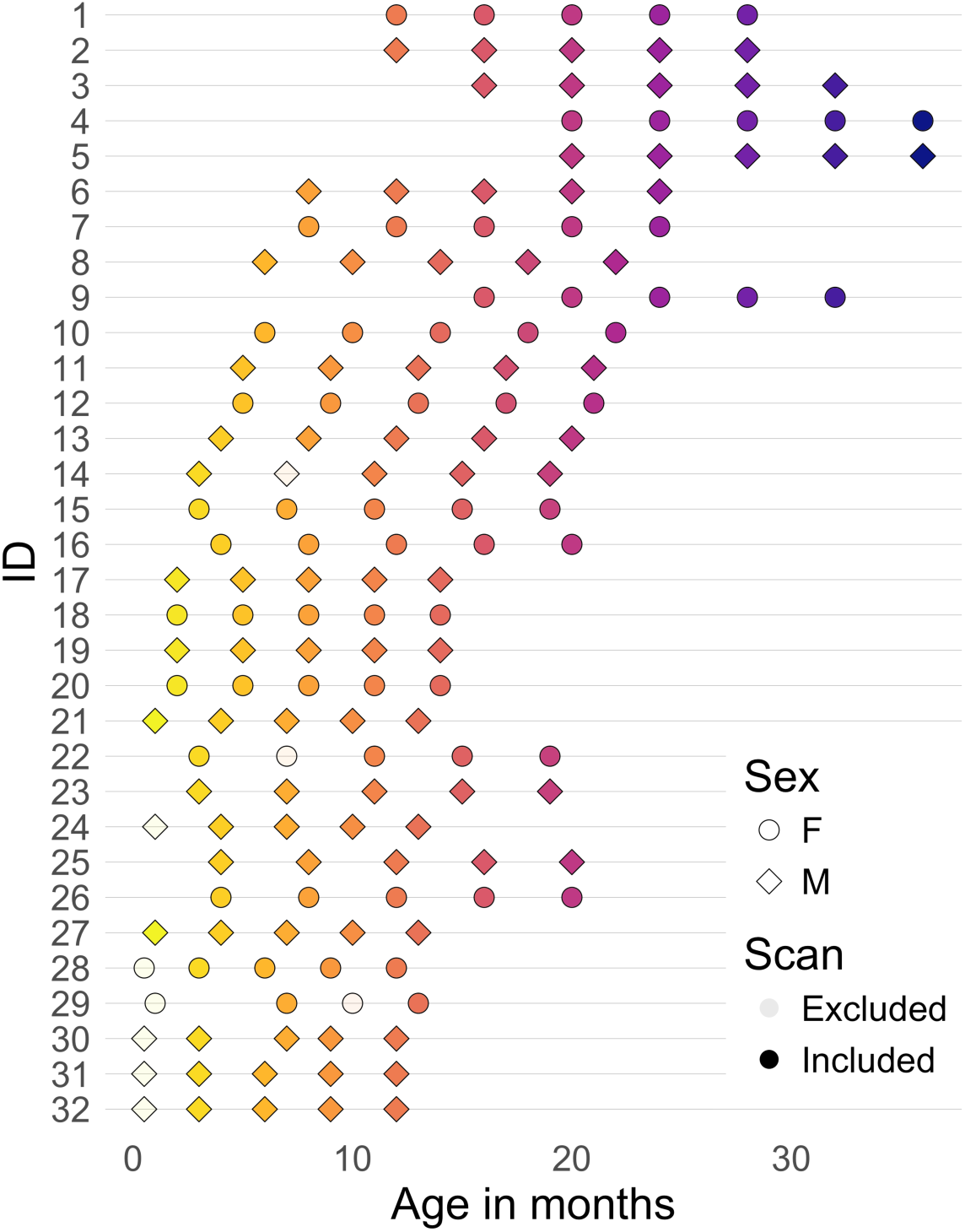
Longitudinal MRI scanning schedule. Up to five MRI scans were acquired from each macaque. Plot shows the age at scan for each macaque (ID). Colour indicates age, transparency indicates timepoints excluded from the analysis, sex is shown by shape.

### Neuroimaging

MRI scans were performed on a 3T scanner (General Electric Medical, Milwaukee WI) using a human 8-channel brain array coil at the University of Wisconsin-Madison. High-resolution 3D T1-weighted imaging was performed using the following parameters: TI = 450 ms, TR = 8.684 ms, TE = 3.652 ms, FOV = 140 × 140 mm, flip angle = 12°, matrix = 256 × 256, thickness = 0.8 mm, gap = −0.4 mm, total time = 10:46 min, providing an effective voxel resolution of 0.55 × 0.55 × 0.8 mm. As part of the scanning protocol, T2-weighted scans and 120-direction diffusion-weighted scans were also acquired (Young et al. 2017). Macaques were anaesthetised for the duration of the scan.

### Image processing

The image processing pipeline is shown in Fig 2. Raw T1 images were initially denoised using a non-local means filter (Manjón et al. 2012) and aligned using a rigid registration to the NIMH Macaque Template (NMT)(Seidlitz et al. 2018).

**Figure 2:**
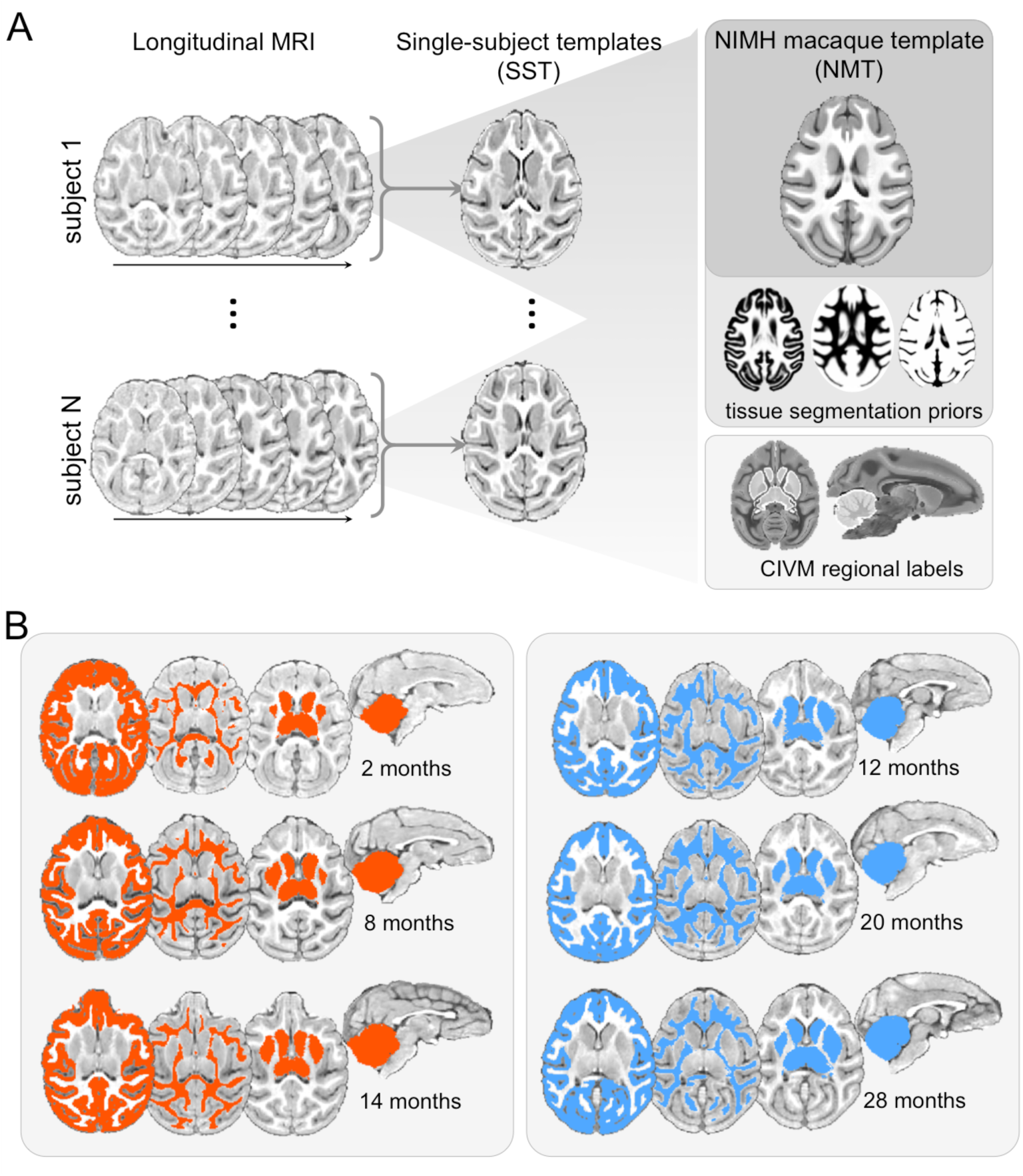
Image processing pipeline and example tissue segmentations. A. Longitudinal MRI scans were aligned to construct single-subject templates (SST) that were, in turn, aligned to a macaque template image. SST images were segmented using tissue priors from the NMT template, producing subject-specific tissue priors to initialise segmentation in each MRI image. Subcortical and cerebellar regions were labelled using the CIVM atlas. B. Tissue segmentations are shown at three timepoints for two macaques (orange and blue).

To perform brain extraction, T1-weighted MRI were corrected for bias field inhomogeneities and aligned to the template image using a diffeomorphic nonlinear registration in ***ANTs*** (Avants et al. 2008, 2011; Tustison et al. 2010). Images were brain extracted using a probabilistic tissue segmentation (***Atropos*** in ***ANTs;*** Avants et al. 2011) initialised by the NMT tissue priors. Brain masks were visually checked and edited where necessary to ensure non-brain tissue was removed from individual brain masks.

To account for the longitudinal structure of the dataset, single subject templates (SST) were constructed by iterative nonlinear alignment and averaging of all available timepoints for a given subject (Avants et al. 2011; Reuter et al. 2012), resulting in 32 SST (Figure 2A). Nonlinear transformations were calculated between each subject’s individual timepoints and the corresponding SST, and between each SST and the NMT. Each SST was segmented into three tissue classes (grey matter: GM; white matter: WM, and cerebrospinal fluid (CSF) with ***Atropos*** using the spatial tissue priors of the NMT. Individual tissue priors were created by smoothing the SST tissue segmentations and used to initialise tissue segmentation of each individual’s timepoint image (Figure 2B).

Subcortical structures were labelled using regional tissue labels derived from the Center for In Vivo Microscopy (CIVM) macaque atlas, consisting of 242 regional parcels hierarchically organised according to the developmental ontology of the mammalian brain (Puelles et al. 2013; Calabrese et al. 2015). Labels were combined hierarchically into a subcortical grey matter mask (comprising the thalamus, pallidum, caudate, putamen, amygdala and hippocampus) and a cerebellar mask. Regional masks were then propagated to individual scans via the concatenated warps computed between the NMT and individual SSTs (Fig 2A), and between each SST and each individual timepoint. Voxels contained by each mask, but not classified as CSF, were classified as subcortical or cerebellar voxels respectively. This resulted in WM, GM, subcortical GM and cerebellar masks for each T1 image (Fig 2B).

A visual quality check was performed for all images, with gross misclassification errors edited manually where necessary. The isointensity of different tissue classes in MRI of the early postnatal brain (Holland et al. 1986; Dietrich et al. 1988) can render tissue segmentation problematic (Weisenfeld and Warfield 2009) and prone to significant error. All T1 weighted scans acquired earlier than 1 month (n=4) were excluded from further analysis due to largely failed tissue segmentation. A further 5 timepoints were removed due to failed image registrations or tissue segmentations (Figure 1).

Thus, for each timepoint, we calculated volumes for the whole brain, cortical and subcortical grey matter, cerebral white matter and cerebellum for statistical analysis.

### Statistical analysis

We use Generalised Additive Models (GAMs) to model the relationship between brain tissue volume and age. Specifically, we can model the relationship between tissue volume, *y* and age for *n* subjects as:

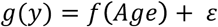

Where *f* (·)represents an unknown smooth function of age,*g*(·) represents a link function to map the distribution of the predicted values to that of the response and ε is an *n* × 1 vector of errors, or residuals, assumed to be 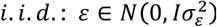. In the case that *y* is normally distributed, *g*(*y*) = *y*.

This model can be extended to include multiple linear and nonlinear predictor variables, and when error is correlated across samples (i.e.: with repeated measures of the same individual), by incorporating random effects to model the covariance between within-subject observations:

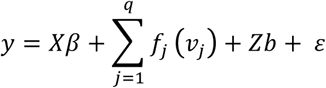

Where *X* is an *m*-observation ×*p* matrix modelling *p* linear fixed effects,{*v*_1_,…,*v*_*q*_} is a set of *q* nonlinear predictors and *Z* is an *m* × (*n* · *r*) design matrix modelling *r* random effects across *n* subjects. The linear parameters, *β*, and smooth functions,*f*_1_…_*p*_(·), can then be estimated along with predictions of the random effects, *b*.

In this study, we used the ***mgcv*** package (v1.8-22) implemented in ***R*** (v3.3.1) to model longitudinal tissue volume trajectories with GAMs. Smooth functions were estimated using penalised thin plate regression splines with automatic smoothness estimation (Wood 2003, 2011).

To account for the longitudinal nature of the data, we included random effects to model subject-specific deviations from the group intercept or age relationship (random intercepts and slopes, respectively). We fit two mixed effects models for each tissue volume with fixed linear effects of sex and a smooth function of age as well as either random intercepts (***GAM1***), or both random intercepts and slopes (***GAM2***). For all tissue compartments (GM, WM, scGM and cerebellum), we also included a fixed linear effect of intracranial volume. For each tissue volume, we fit a separate model with the additional inclusion of an age:sex interaction effect for comparison. We compared the nonparametric GAM to a baseline linear mixed effects model with random intercepts using the ***lme4*** package (v1.1-15). Models were fit using Maximum Likelihood and compared with the Akaike Information Criterion (AIC).

To test the contribution of each term to the final model for each tissue compartment, we fit a GAM including all terms and random effects and calculated the difference in explained variance excluding terms one at a time. Removal of terms that are important to the model fit will result in a larger reduction in variance explained. Wald tests were performed to test the significance of each parametric and smooth term’s contribution to the model.

## Results

### Intracranial volume

Longitudinal trajectories for intracranial volume are shown in Figure 3. On average, ICV followed a nonlinear trajectory over time with an initial increase from 81.2 [95% C.I.: 78.4,84.0] ml at 1 month to 95.6 [93.3,97.7] ml at 5 months, a 17.8% increase in volume. This was followed by a small 2-3% decrease to approximately 92.7 [90.4,95.0] ml at 9 months and a slow, relatively stable increase of around 0.35ml (0.37%) per month to reach 102.3 [98.4,106.3] ml at 36 months (Fig 3)

**Figure 3:**
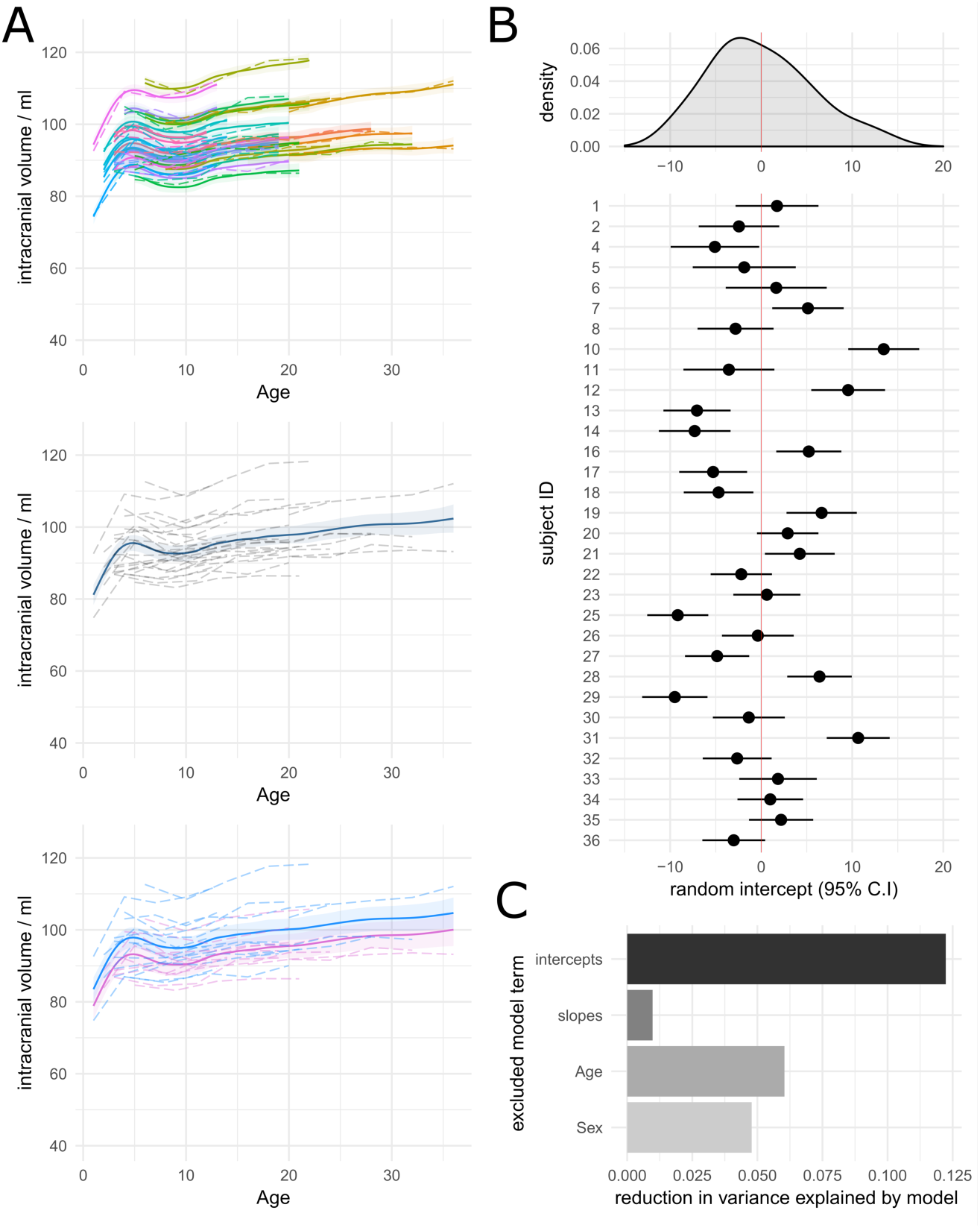
Modelling longitudinal change in intracranial volume. A. Modelled longitudinal trajectories are shown for all individuals (top row), the whole group (middle) and for males (blue) and females (pink) separately (bottom). Raw data are shown with dotted lines. Solid lines indicate predicted volume with 95% confidence intervals shaded. **B.** The distribution of random intercepts (histogram, top) and prediction of individual random effects with 95% C.I. are shown. **C.** Relative importance of models terms in the full model.

A summary of model fits is shown in Table 1. Overall, the nonlinear models of ICV change produced a better fit of the data compared to a linear model. A Chi-squared test on the maximum-likelihood scores of the GAM models revealed a marginal improvement in the model fit when including both random intercepts and slopes (ML scores: ***GAM1***=372.46, ***GAM2***=369.41; p=0.014). However, the inclusion of random slopes only increased the variance explained by the model by ∼1% (Table 3, F_14.8,31_=221.9, p=0.02) and variation in the relationship between age and ICV was relatively low across subjects (Fig S1). Fixed effects parameters for the best-fit model (including intercepts and slopes) are shown in Table 2.

**Table 1:**
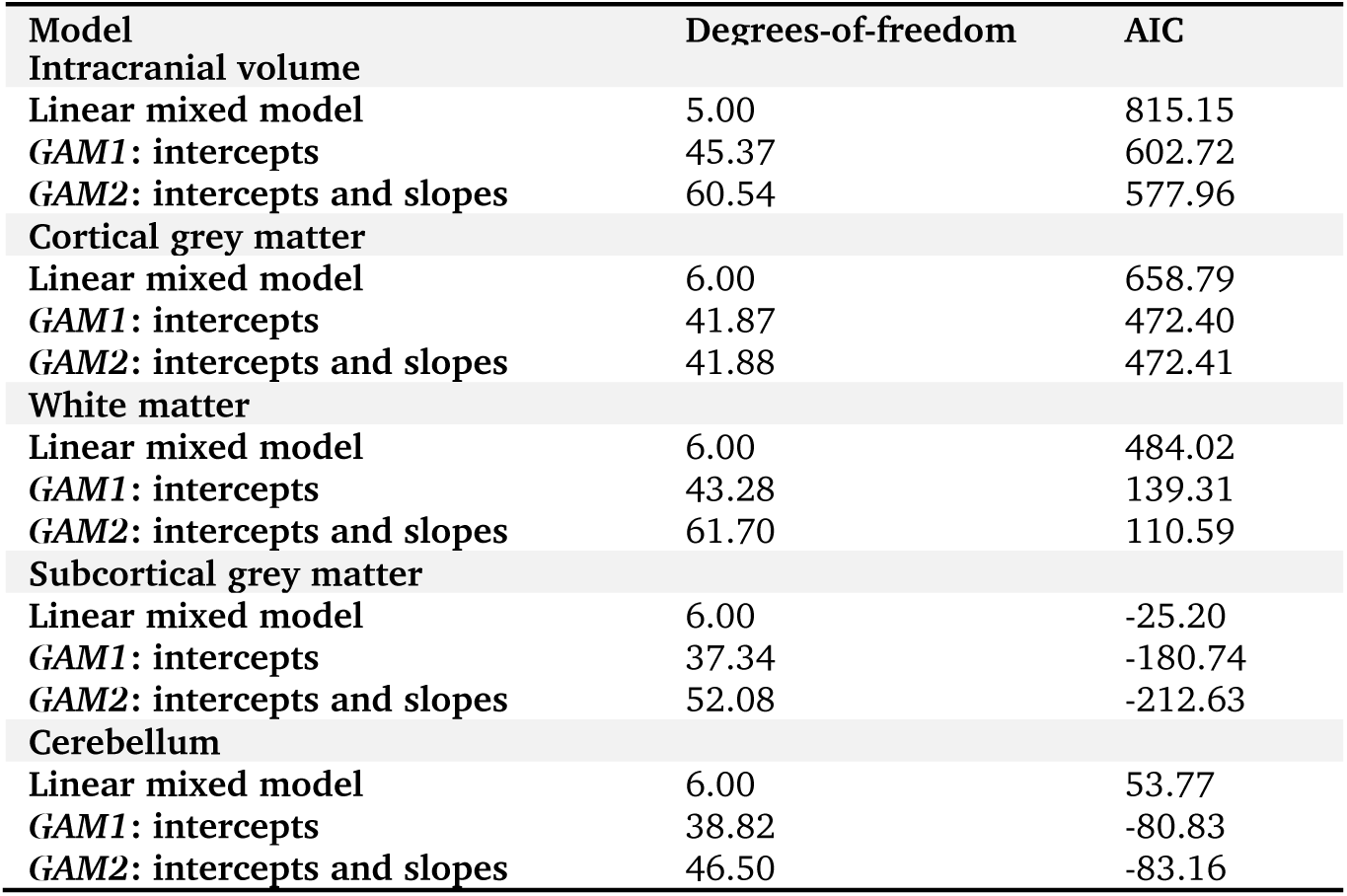
Model goodness-of-fit summaries.

| Model | Degrees-of-freedom | AIC |
| --- | --- | --- |
| **Intracranial volume** |  |  |
| Linear mixed model | 5.00 | 815.15 |
| **GAM1: intercepts** | 45.37 | 602.72 |
| **GAM2: intercepts and slopes** | 60.54 | 577.96 |
| **Cortical grey matter** |  |  |
| Linear mixed model | 6.00 | 658.79 |
| **GAM1: intercepts** | 41.87 | 472.40 |
| **GAM2: intercepts and slopes** | 41.88 | 472.41 |
| **White matter** |  |  |
| Linear mixed model | 6.00 | 484.02 |
| **GAM1: intercepts** | 43.28 | 139.31 |
| **GAM2: intercepts and slopes** | 61.70 | 110.59 |
| **Subcortical grey matter** |  |  |
| Linear mixed model | 6.00 | -25.20 |
| **GAM1: intercepts** | 37.34 | -180.74 |
| **GAM2: intercepts and slopes** | 52.08 | -212.63 |
| **Cerebellum** |  |  |
| Linear mixed model | 6.00 | 53.77 |
| **GAM1: intercepts** | 38.82 | -80.83 |
| **GAM2: intercepts and slopes** | 46.50 | -83.16 |

**Table 2:** Parameter estimates for fixed linear effects in best fit models.

| Term | Estimate | 95% C.I.<br>lower | upper | Std. est. | 95% C.I.<br>lower | upper | p |
| --- | --- | --- | --- | --- | --- | --- | --- |
| **Intracranial volume** |  |  |  |  |  |  |  |
| Intercept | 92.99 | 89.83 | 96.15 | - | - | - | - |
| Male | 4.64 | 0.49 | 8.80 | 0.312 | 0.03 | 0.59 | 0.03 |
| **Cortical grey matter** |  |  |  |  |  |  |  |
| Intercept | -3.41 | -8.98 | 2.17 | - | - | - | - |
| Male | -0.62 | -1.54 | 0.29 | -0.07 | -0.18 | 0.04 | 0.184 |
| ICV | 0.54 | 0.48 | 0.60 | 0.95 | 0.85 | 1.06 | <0.001 |
| **White matter** |  |  |  |  |  |  |  |
| Intercept | -2.99 | -5.86 | -0.12 | - | - | - | - |
| Male | 0.55 | -0.21 | 1.30 | 0.08 | -0.03 | 0.19 | 0.16 |
| ICV | 0.20 | 0.17 | 0.23 | 0.42 | 0.36 | 0.49 | <0.001 |
| **Subcortical grey matter** |  |  |  |  |  |  |  |
| Intercept | 1.26 | 0.58 | 1.93 | - | - | - | - |
| Male | 0.02 | -0.13 | 0.16 | 0.01 | -0.09 | 0.11 | 0.84 |
| ICV | 0.06 | 0.05 | 0.07 | 0.60 | 0.53 | 0.67 | <0.001 |
| **Cerebellum** |  |  |  |  |  |  |  |
| Intercept | 1.61 | 0.59 | 2.638 | - | - | - | - |
| Male | 0.05 | -0.17 | 0.27 | 0.04 | -0.11 | 0.73 | 0.63 |
| ICV | 0.06 | 0.05 | 0.07 | 0.62 | 0.51 | 0.73 | <0.001 |

**Table 3:**
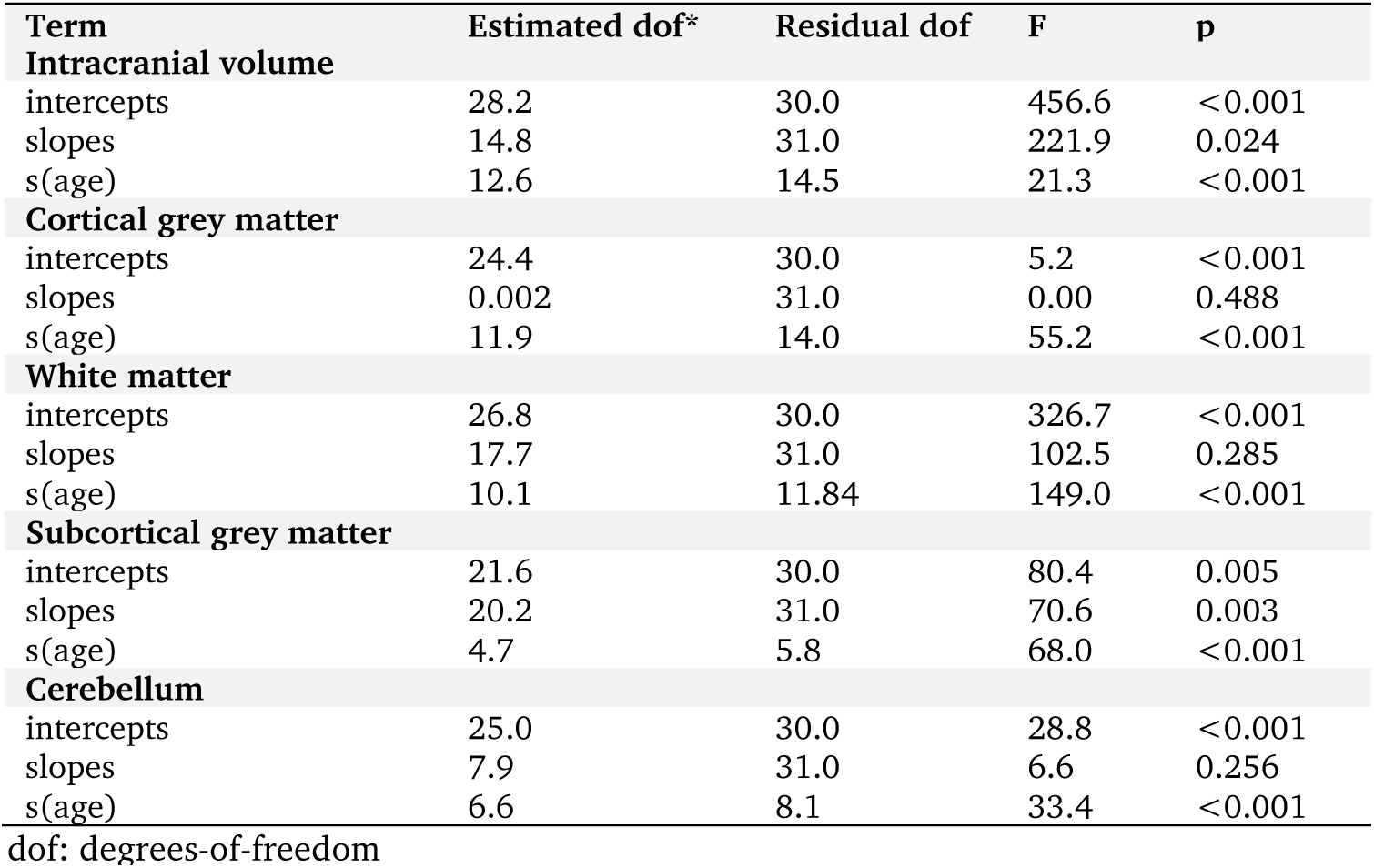
Significance of all smooth terms and random effects in mixed effects models.

The inclusion of random intercepts resulted in a 12.2% increase in variance explained over other terms (Fig 3C). Modelling age-related change in ICV also explained a significant amount of variance in the data (∼6%).

The distribution of predicted random intercepts for each subject are shown in Fig 3B, and for random slopes in Fig S1. A large variation in head size was observed across the cohort. The group mean intercept [95% C.I] was 92.99ml [89.83, 96.15](Table 2), in this case representing the mean ICV in females with individual ICV varying around this by ±15-20ml. This variation is apparent in Fig 3A (top row) where individual trajectories are overlaid on the volumetric data.

The inclusion of sex as a factor improved the variance explained by the model by around 4.8% (Fig 3C). ICV was significantly larger in males than females by an average 4.64ml [0.49,8.80] (p=0.031). Average male and female trajectories are shown in Fig 3A (bottom row). Adding an additional model term to test whether the smooth function of age varied between sexes did not improve model fit (AIC with interaction effect: 585.3).

### Cortical grey matter volume

On average, absolute CGM volume increased from 47.0 [44.4, 49.6] ml at 1 month to a maximum of 51.5 [49.6, 53.4] ml at 4 months before decreasing to 45.8 [42.8, 48.8] ml at 36 months with CGM volume moderately greater in males than females (mean=2.35ml [0.21, 4.48], p=0.03; Fig S2)

As expected, CGM was largely dependent on ICV (Fig 4A, bottom; Table 2). After correcting for linear scaling effects of ICV, cortical grey matter volume followed a nonlinear trajectory between 1 and 36 months. On average, cortical grey matter volume decreased, as a proportion of ICV, over the whole period (Fig 4A, top). This trajectory was characterised by an initial period of rapid decline between 1 and 8 months from 59.6% of ICV to 49.2% and a longer period of slower decrease, reaching an apparent plateau of around 45% by 30 months.

**Figure 4.**
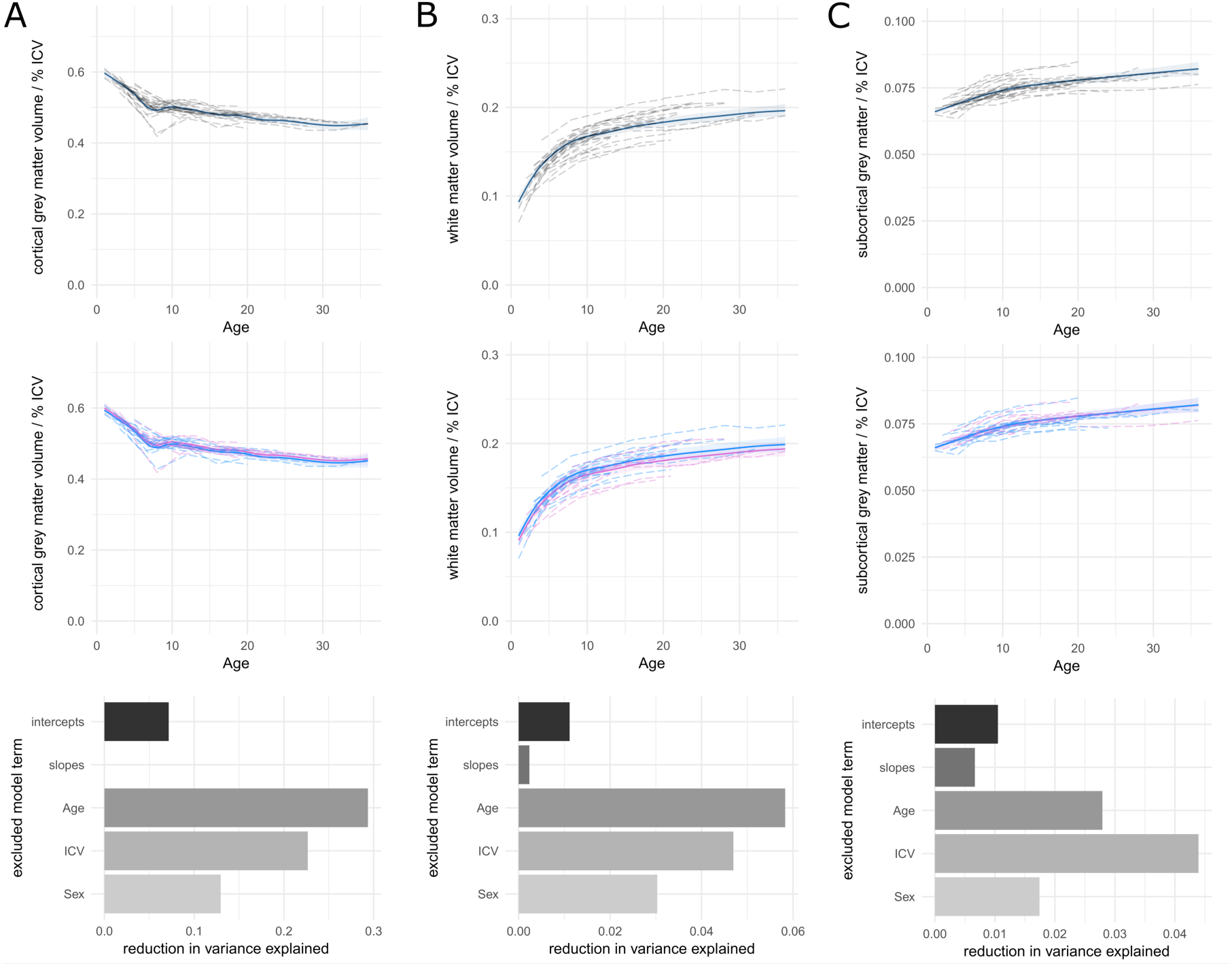
Longitudinal trajectories of tissue volume. Group average (top row) and male and female (middle row) trajectories are visualised as a proportion of ICV for cortical grey matter (A), white matter (B) and subcortical grey matter (C). Raw data are shown with dotted lines, solid lines represent model fit with 95% C.I. shaded. Bottom row shows the reduction in variance explained when excluded model terms one-by-one from the full model fit. Trajectories for absolute volume are shown in Fig S2.

After accounting for variance due to individual differences in ICV, the inclusion of random intercepts significantly increased variance explained by the model (Table 3; F_24.4,30.0_=456.6, p<0.001) but the inclusion of random slopes did not (p=0.488). The best-fit model therefore included random intercepts only (Table 1). The inclusion of age (29%) or intracranial volume (23%) terms improved model fit significantly (Tables 3; Fig 4A, bottom) but there was no significant difference in relative cortical grey matter volume between sexes after accounting for ICV (Table 2; Fig 4A, middle; p=0.18).

### White matter volume

Absolute WM volume increased from 6.6 [5.8,7.4] ml at 1 month to a maximum of 20.4 [19.3,21.5] ml at 36 months (Fig S2). On average, total WMV was greater in males than females by 1.4ml ([0.18, 2.57], p=0.026), although this difference was not significant when including ICV in the full model (Table 2, Fig 4B, middle).

White matter volume increased as a proportion of ICV over time from 9.3% to 19.6%, most rapidly over the first 7-8 months (Fig 4B). Goodness-of-fit estimates are shown in Table 1. The inclusion of both random intercepts and random slopes led to a marginally better model fit over the inclusion of random intercepts alone (𝒳_2_= 2.53, *p* = 0.025). However, the variation in white matter change over time did not vary greatly over subjects and inclusion of this term did not significantly increase the variance explained by the model (Table 3; Fig 4B, bottom). In contrast, age and ICV accounted for relatively large improvements in variance explained by the model over other terms.

### Subcortical grey matter volume

Subcortical grey matter volume increased from 5.6 [5.3,5.8] ml to 8.3 [7.9,8.7] ml between 1 and 36 months (Fig S2). There was no significant difference between males and females in absolute volume (*β* =0.29 [-0.03,0.61], p=0.08). As a proportion of ICV, subcortical grey matter volume increased from 6.6% to 8.2% (Fig 4C).

The relative importance of each model term is shown in Fig 4C (bottom row). The best fit model included both random intercepts and slopes (Table 1; 𝒳_2_ = 8.15, *p* < 0.001) and the inclusion of both random intercept and random slope terms independently increased the variance explained by the model (Table 3, both p<0.01). Figure 5 shows individual variation in predicted subcortical volume across the full observation window for all subjects after accounting for the fixed effects of ICV and sex.

**Figure 5:**
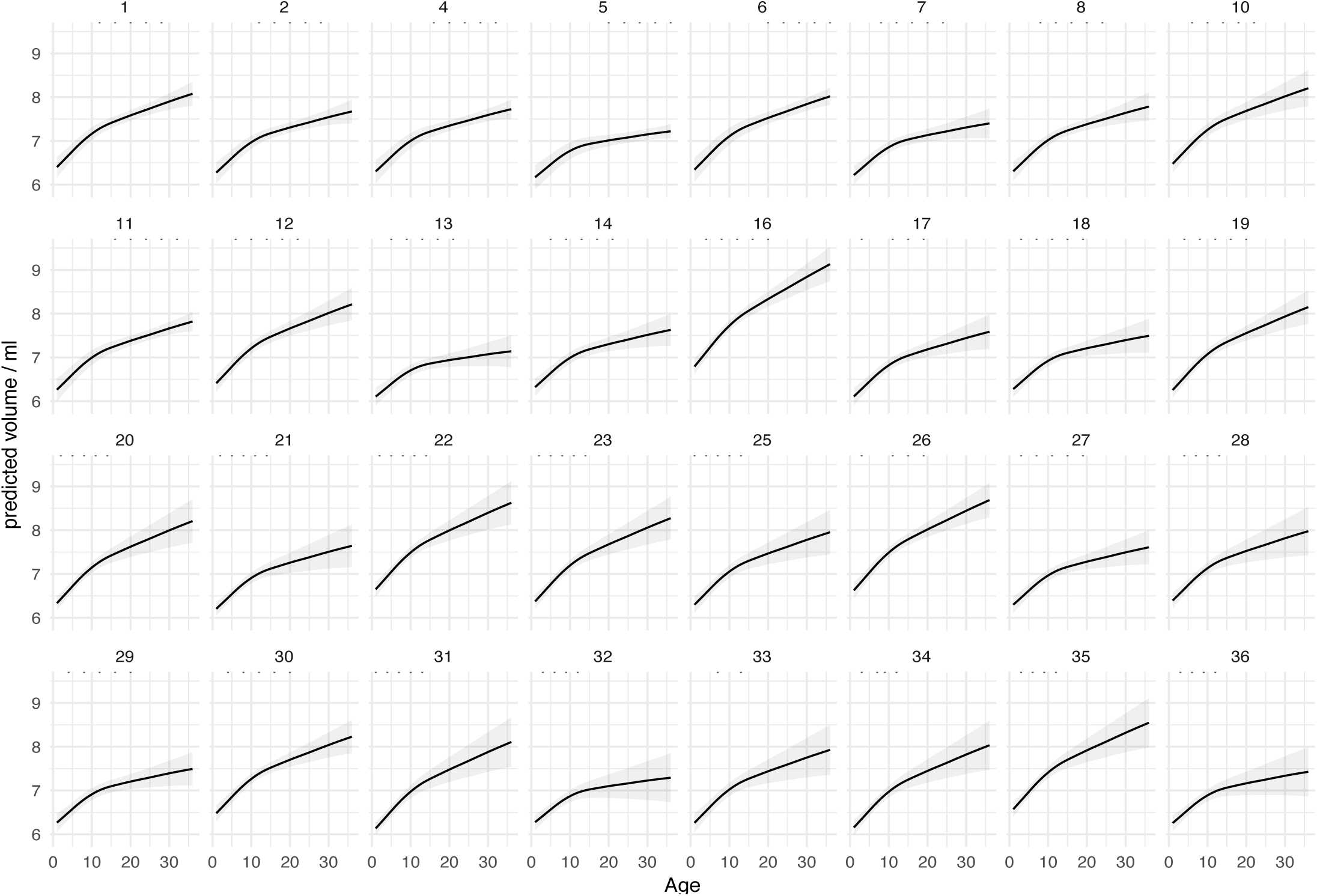
Individual variation in subcortical grey matter volume. Predicted trajectories of subcortical grey matter volume for every subject with 95% C.I. shaded accounting for differences in ICV and sex. Dots below each subject ID show the age at which individual MRI scans were acquired.

### Cerebellar volume

Cerebellar volume increased from 5.56 [5.3,5.8] ml (6.8% of ICV) at 1 month to 8.6 [8.3,8.9] m (8.9%) at 36 months (Fig S2). Overall the inclusion of ICV (Table 2), age and random intercepts (Table 3) significantly increased explained variance (Fig S3; all p<0.001), with the best fit model including random intercepts only (Table 1; *𝒳*_2_= 0.67, p=0.25). There was a significant effect of sex on absolute cerebellar volume (*β* =0.40 [0.09,0.72], p=0.014), but not after including ICV in the full model (Table 2, p=0.63).

### Relating individual variations in tissue volume to birth weight

Birth weight was available for 31 of 32 macaques (mean = 535.7g, range: 425-672g). We regressed individual predictions of random model effects (intercepts, and slopes when included in the model) onto birth weight for each tissue compartment. We found that lower birth weight was associated with lower subcortical grey matter volume after accounting for ICV. Birth weight explained 23% of the variance in subcortical volume model intercepts (R^2^=0.23, F=8.62, p=0.006, p<0.05 after correction for multiple comparisons). Individual variation in volume did not correlate with birth weight for ICV (p=0.39), WM (p=0.49) or cerebellum (p=0.20). There was a small association between lower birth weight and higher relative cortical grey matter (R^2^=0.15, F=4.92, p=0.035), though this did not survive correction for multiple comparisons. There were no associations between birth weight and rate of change of volume over time (random slopes) for ICV (p=0.068) or subcortical grey matter (p=0.324).

## Discussion

In this paper, we present data-driven, nonlinear models of postnatal brain growth in the macaque across an age range broadly equivalent to 3 months to 9 years in humans. Intracranial volume significantly increased between 1 and 36 months. After accounting for changes in ICV over time, cortical grey matter decreased overall, whereas white matter, subcortical grey matter and cerebellar volume all increased in volume. We also found that absolute intracranial volume varied significantly between individuals. After accounting for inter-individual differences in ICV, rate of change in cortical grey matter, white matter and cerebellar volume were relatively stable within the group, whereas rate of change of subcortical grey matter volume varied significantly across subjects. Lower birth weight was not associated with differences in ICV, but was significantly associated with lower subcortical grey matter volume and moderately larger cortical grey matter volumes.

Intracranial volume, encompassing whole brain tissue as well as the ventricular system, increased by 25% between 1 and 36 months of age along a relatively stable, nonlinear trajectory with two notable deviations: a period of relatively faster volume increase between 1 and 5 months followed by a period of slowed or decreasing volume between 5 and 9 months. This stalling in brain growth has been observed previously in the macaque brain. Scott et al. observed a transient plateau of brain growth between 26 and 39 weeks (approximately 6 to 9 months) in two smaller, independent cohorts (N=24 and 21) with MRI acquired on different scanners (Scott et al. 2016). This was characterised by a lack of growth in male and negative growth in the female brain, along with a slowing in somatic growth during the same period. In this study, we observe a similar plateau, although we do not find significant evidence for different trajectories in male and female brain. In a smaller longitudinal cohort (n=7), a similar pattern was reported, with decreasing brain volume observed in 4 out of 7 monkeys between 5 and 8 months that resulted in a group average decrease of 3% (Malkova et al. 2006). Here, we confirm these findings in a larger, developmental cohort.

While many neurodevelopmental processes are conserved across species, defining equivalent stages across species remains a complex task (Workman et al. 2013). Previous studies have approximated human age as equivalent to three times macaque age based on a comparison of lifespan and the age of pubertal maturation across species (Tigges et al. 1988; Chen et al. 2013). Based on this estimate, the period of slowed brain growth should fall between approximately 1 and 2 years of age in humans. Although few, longitudinal studies of infant brain development during this period have found that brain volume increases rapidly between birth and one year of age, doubling in size to reach 72% of adult volume. Between 1 and 2 years, growth rate decreases significantly with volume increasing by 15% over the second year (Knickmeyer et al. 2008). In a seminal study of brain weight, Dekaban and Sadowsky reported a dramatic slowing of growth rates by 3 years of age (Dekaban and Sadowsky 1978). In both studies, these patterns reflect changes in total brain tissue volume (i.e.: parenchymal tissue, not including the ventricular system) and over the same period of time, the volume of the lateral ventricles was found to decrease by around 8% across the group (24% in males) (Knickmeyer et al. 2008). While this suggests that the observed decreases in ICV may be due to changes in ventricular volume rather than brain tissue, later work in a larger cohort estimated ventricular volume to increase around 18% between 1 and 2 years in humans (Knickmeyer et al. 2008; Gilmore et al. 2012). Alternative explanations may include a period of imbalance between progressive (axonal outgrowth, dendritic arborisation) and regressive (synaptic pruning) neurodevelopmental processes that dramatically slow brain growth, or that decreases in brain volume are a direct consequence of decreased somatic growth (Passingham 1985; Bourgeois and Rakic 1993; Bourgeois et al. 1994; Scott et al. 2016).

From 10 months onwards, ICV was observed to increase slowly, gaining on average an additional 10% volume between 10 and 36 months (Fig 3). In humans, early cross-sectional studies suggested the ICV peaks at around 10 years, approximately equivalent to 40 months in the macaque (Pfefferbaum et al. 1994). Longitudinal studies have since confirmed that ICV continues to increase steadily from mid-childhood before stabilising in adolescence, with an annual increase of around 1% reported (Giedd et al. 1999; Mills et al. 2016).

Using mixed effects models, we found significant inter-individual variation in ICV across the group. Brain volume at 1 month was predicted to vary by ±15-20 ml around the group average. This represents substantial individual variation given that the average difference between male and female brains at the same age was estimated to be under 5ml. No other differences were found between sexes across the other tissue compartments, after correcting for differences in ICV. This pattern highlights the importance of considering sample variation when comparing two or more populations. When considering neuroanatomical sex differences in humans, volumetric distributions are largely overlapping with relatively small mean difference (Ritchie et al. 2018). Similarly overlapping distributions have been reported in clinical populations including ADHD, autism and schizophrenia (Erp et al. 2016; Hoogman et al. 2017; van Rooij et al. 2018), highlighting the importance of longitudinal modelling to extract within-individual differences relative to the sample mean.

By comparison, cortical grey matter volume remained relatively stable over time. In absolute terms, CGM increased initially, gaining 10% volume between 1 and 4 months, before decreasing to plateau by around 10 months. In humans, Gilmore et al. reported an increase in total cortical grey matter of 107% between birth and 12 months (Gilmore et al. 2012) While significantly greater than the changes reported in macaques, we speculate that the period of growth observed here represents the end of a period of rapid change. Histological evidence shows that thickness of the striate cortex is maximal at around 3 months of age, increasing by around 32% from thickness at birth (Bourgeois and Rakic 1993). The newborn macaque brain is approximately 60% of adult volume, compared to 25% in humans (Sacher and Staffeldt 1974; Malkova et al. 2006) suggesting a period of neoteny that is relatively longer in humans. As such, it is likely that a significant increase in cortical grey matter is evident prior to one month in the macaques. However, segmentation of tissue compartments on MRI becomes difficult at younger ages due to the lack of contrast between grey and white matter on T1 and T2-weighted images (Holland et al. 1986; Dietrich et al. 1988). Tissue segmentations at this point rely heavily on manual tracing or specialised algorithms (de Macedo Rodrigues et al. 2015; Moeskops and Pluim 2017). In this study, we excluded scans acquired at less than one month due to errors in segmentation thus limiting our ability to define cortical development during this stage of development. Previous longitudinal studies in the macaque did not report grey matter volumes, instead estimating lobar volume inclusive of both white and grey matter (Scott et al. 2016), or manually traced volumes of white matter tissue alone (Malkova et al. 2006). In a cross-sectional study of macaques aged 10 to 64 months, total cortical grey matter remained stable across the observation window, as reported here (Knickmeyer et al. 2010).

While extant longitudinal data in humans is relatively sparse between 2 and 5 years of age, cortical grey matter is highest in childhood and appears to decline from age 5 onwards (Lebel and Beaulieu 2011; Mills et al. 2016). In our study, absolute cortical grey matter did not decline significantly with age after 10 months. However, cortical grey matter was significantly dependent on changes in intracranial volume and when ICV was included in the model, the effect of age on cortical grey matter followed a clear downward trajectory, mirroring that seen in humans when changes in ICV is taken into account (Mills et al. 2016). When visualised as a proportion of ICV, CGM decreased from an average of 60% to 45% (Fig 4). After accounting for the substantial individual variation in ICV, the overall amount of cortical grey matter continued to vary across subjects, but the rate of change of cortical grey matter over time was relatively conserved. In humans, the rate of change in cortical grey matter volume varies significantly across cortical regions with developmental changes in higher-order association cortex occurring later than in sensorimotor regions (Giedd et al. 1999; Sowell et al. 2004; Gilmore et al. 2012) and similar patterns have been reported in macaques, even when total grey matter remains stable (Knickmeyer et al. 2010; Scott et al. 2016). This suggests that the amount of individual variability in cortical grey matter change may vary over regions and over time, and some aspect of this subject-level variation may be lost when considering the cortex as a whole.

White matter volume increased significantly over time both in absolute terms and as a proportion of ICV, with rapid expansion in the first eight months and continued growth up to 36 months, confirming previous reports (Malkova et al. 2006). In humans, white matter continues to increase into mid-to-late adolescence (Lebel and Beaulieu 2011; Mills et al. 2016), and in macaques WM appears to peak post-puberty in young adulthood at approximately 4 years (Knickmeyer et al. 2010). This volumetric increase likely reflects the extent of ongoing myelination processes within the cerebral white matter. In humans, white matter myelination follows a protracted developmental course, extending into the second decade of life in humans (Yakovlev and Lecours 1967). Axonal myelination in non-human primates is more widespread at birth compared to humans and develops along a similar, though less prolonged, trajectory until adolescence (Malkova et al. 2006; Miller et al. 2012). After accounting for variance due to ICV, we found that individual variation in white matter volume remained but, as with the cortical grey matter, rate of change of WM volume was relatively stable across subjects, suggesting that the neurobiological processes governing WM growth over time were relatively well-preserved across subjects.

In contrast, we found that subcortical grey matter exhibited significant inter-individual variation in both volume and rate of change over time, even after accounting for individual differences in ICV. Growth over time was, in general, less dramatic, increasing from 6.6% of ICV to 8.2% by 36 months, with two distinct phases of faster and slower growth up to, and after, 10 months, respectively. Cerebellar volume followed a similar trajectory to the subcortical grey matter, increasing rapidly over the first 8-9 months, however rate of change of cerebellar volume did not vary significantly across subjects. In humans, change in cerebellar and subcortical grey volume are strongly correlated in the second year of life (Knickmeyer et al. 2008), with cerebellar volume increasing until around 12-15 years (Tiemeier et al. 2010).

In humans, subcortical structures increase by around 106% over the first year of life, with rate of hippocampal growth notably slower (Gilmore et al. 2012). In later childhood, the thalamus, pallidum and hippocampus increase in volume into adolescence before decreasing whereas caudate and putamen volumes decrease from around the age of 5 onwards (Herting et al. 2018). In macaques, manual tracing of the hippocampus reveal a similar trajectory to that described in humans, peaking by 3 years of age before declining slightly (Payne et al. 2010; Hunsaker et al. 2014). In contrast, using cross-sectional data, Knickmeyer described linear increases of caudate, putamen and hippocampal volume between 10 and 64 months in the macaque (Knickmeyer et al. 2010). In this cohort, the variation in development of total subcortical grey matter over time is shown in Figure 5 after accounting for the effects of ICV. All subjects followed the same trajectory, with some predicted larger subcortical volumes overall (e.g.: 22), and some with markedly steeper increases in volume over time (e.g.: 16).

We found that lower birth weight was significantly associated with reduced subcortical volume, after accounting for differences in ICV. While birth weight in this cohort was within the normal range for macaques (Fujikura and Niemann 1967), variation in birth weight can be considered a sensitive marker of the suitability of the intrauterine environment and low birth weight in humans is associated with poor cognitive performance and adverse cerebral development in infancy and increased risk for neuropsychiatric disorder (Morsing et al. 2011; Sucksdorff et al. 2015; Ball et al. 2017).

In contrast, we found a relatively small association between lower birth weight and increasing cortical grey matter volume (after correcting for ICV) between individuals. In humans, lower birth weight is associated with reductions in relative cortical volume at birth in humans (Tolsa et al. 2004) and, in typically-developing children, increasing weight at birth is associated with specific increases in cortical area (Raznahan et al. 2012; Walhovd et al. 2012). In twins, after correction for intracranial volume, weak, negative associations between gestational age at birth and cortical grey matter volume have been reported (Soelen et al. 2010). Differences in cortical metrics may explain the observed discrepancies in reported here compared to previous human studies. Estimates of cortical volume capture information that relates to both area and thickness, and the contribution of each metric to measured volume may vary over the cortex (Winkler et al. 2018). Further exploration of cortical surface measures in this cohort is a promising avenue for future research.

In humans and macaques, subcortical grey matter appears specifically vulnerable to adverse intrauterine conditions, or early exposure to the extrauterine environment through premature birth. In a longitudinal study of macaque brain development, fetal irradiation during gestation resulted in specific volume reductions of 15-24% were observed in the putamen and cortical grey matter (6-15%) (Aldridge et al. 2012), and in humans, subcortical structures are specifically vulnerable to injury following preterm birth (Boardman et al. 2006; Ball et al. 2012). In typically-developing children, normal variation in birth weight is associated with caudate volume, after correcting for total brain volume (Walhovd et al. 2012), and in a longitudinal study of low birth weight adolescents, lower subcortical grey matter volumes were associated with higher rates of psychiatric symptoms between 15 and 19 years of age (Botellero et al. 2017).

During brain development, there are widespread, dynamic changes in gene expression that peak prenatally and reconfigure after birth (Kang et al. 2011; Bakken et al. 2015). Patterns of regional gene expression in the macaque approximate adult levels by early adolescence, however, changes to gene expression levels begin earlier and change less over time in subcortical structures compared to cortical areas, highlighting the earlier maturation of these structures (Bakken et al. 2015, 2016). This has important implications for developmental variations across subjects. As subcortical development begins earlier, adverse events ***in utero*** may have a long-lasting impact on growth and subsequent cognitive development. For example, expression of autism susceptibility gene networks are enriched in the developing striatum during macaque postnatal development, providing a potential mechanistic link between individual variation in subcortical growth, low birth weight and the emergence of subsequent neurodevelopmental disorders (Lampi et al. 2012; Bakken et al. 2015; van Rooij et al. 2018).

In summary, we examined postnatal brain development in a densely sampled, longitudinal macaque cohort. We found significant inter-individual differences in intracranial volume, and growth of the subcortical grey matter. Future work will focus on the delineation of regional growth trajectories: cortical and subcortical grey matter regions likely have considerable spatial variation in developmental trajectories, the elucidation of which may provide further insight into early primate brain development.

## Acknowledgements

This research was conducted within the Developmental Imaging research group, Murdoch Childrens Research Institute and the Children’s MRI Centre, Royal Children’s Hospital, Melbourne, Victoria. It was supported by the Murdoch Childrens Research Institute, the Royal Children’s Hospital, Department of Paediatrics, The University of Melbourne and the Victorian Government’s Operational Infrastructure Support Program. The project was generously supported by RCH1000, a unique arm of The Royal Children’s Hospital Foundation devoted to raising funds for research at The Royal Children’s Hospital.

We would like to thanks the authors and contributors of the UNC-Wisconsin Rhesus Macaque Neurodevelopment Database. The database was supported by grants from the NIMH (MH901645, MH091645-S1, and MH100031) and the NICHD (HD003352, HD003110, and HD079124).

For the CIVM atlas data, all imaging was performed at the Duke Center for In Vivo Microscopy, an NIH/NIBIB National Biomedical Technology Resource Center (P41 EB015897). Other support was provided by NA-MIC Roadmap for Medical Research (U54 EB005149-01), NIMH (R01 MH091645), NICHD (U54 HD079124), and NIA (K01 AG041211). Brain specimens were provided by the Wisconsin National Primate Research Center (P51 OD011106).

## SUPPLEMENTAL MATERIALS

**Figure S1.**
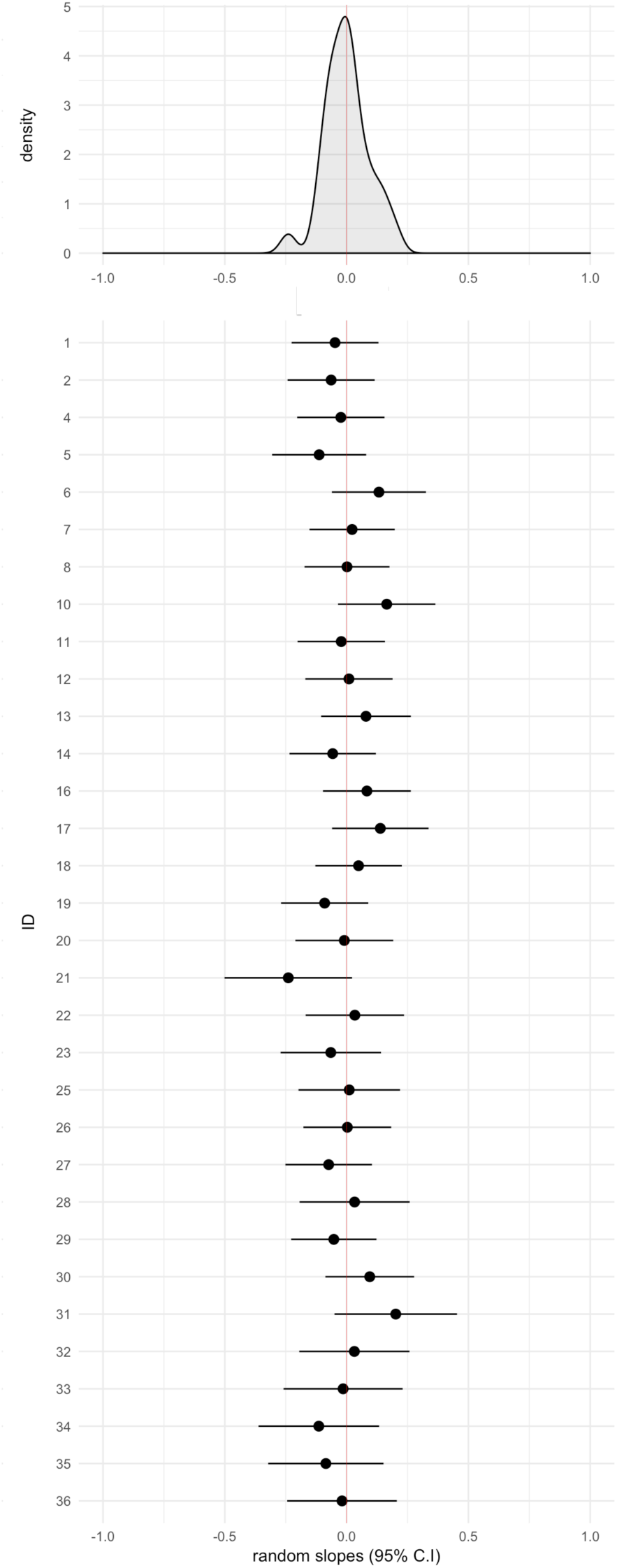
Random slopes for intracranial volume. The distribution of random slopes (histogram, top) and prediction of individual random effects with 95% C.I. are shown for change in intracranial volume over time.

**Figure S2.**
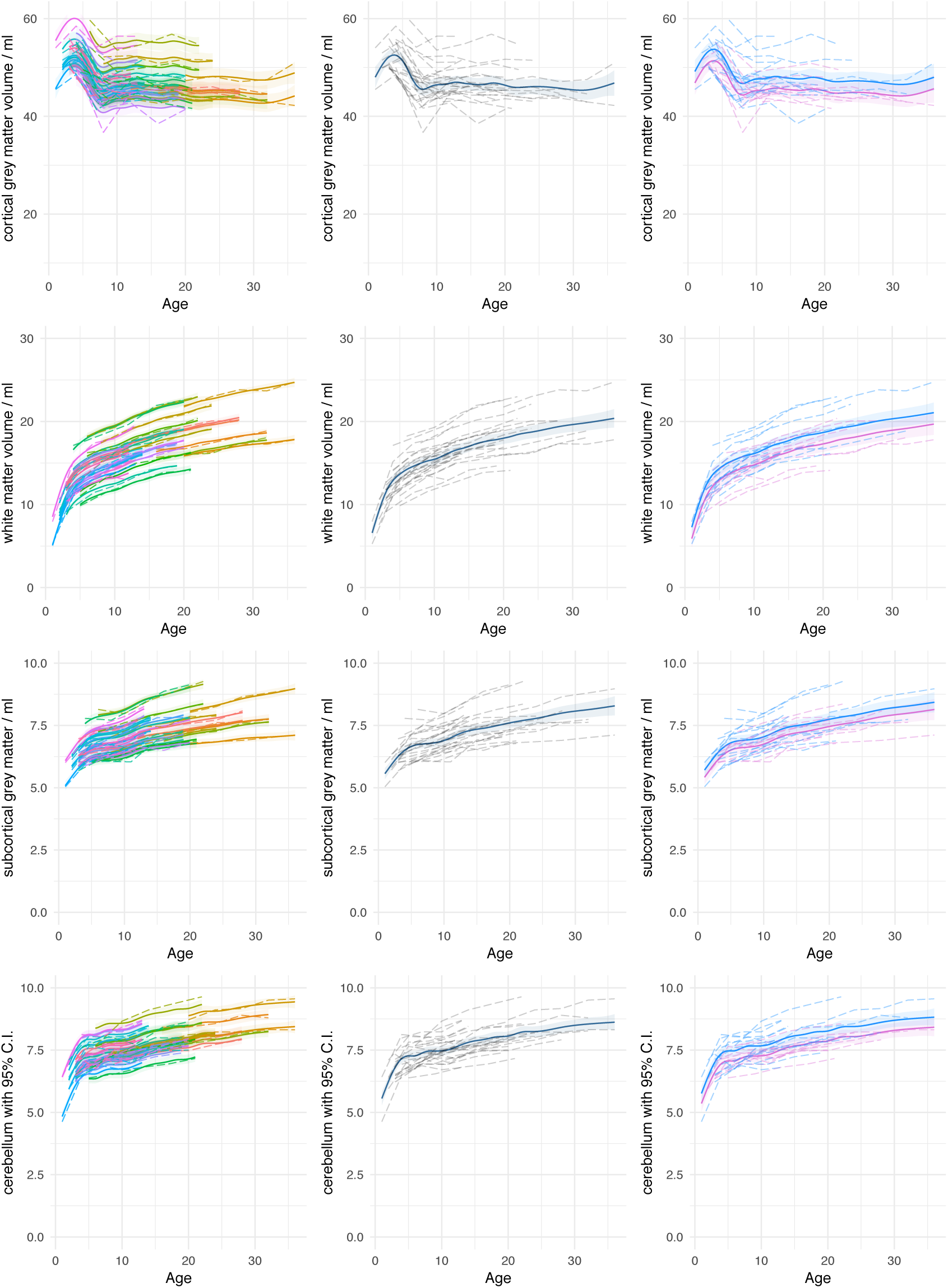
Modelling longitudinal change in absolute tissue volumes volume. Modelled longitudinal trajectories of absolute volumes of cortical grey matter, white matter, subcortical grey matter and the cerebellum are shown for all individuals (left column), the whole group (middle) and for males (blue) and females (pink) separately (right column). Raw data are shown with dotted lines, solid lines represent model fit with 95% C.I. shaded.

**Figure S3.**
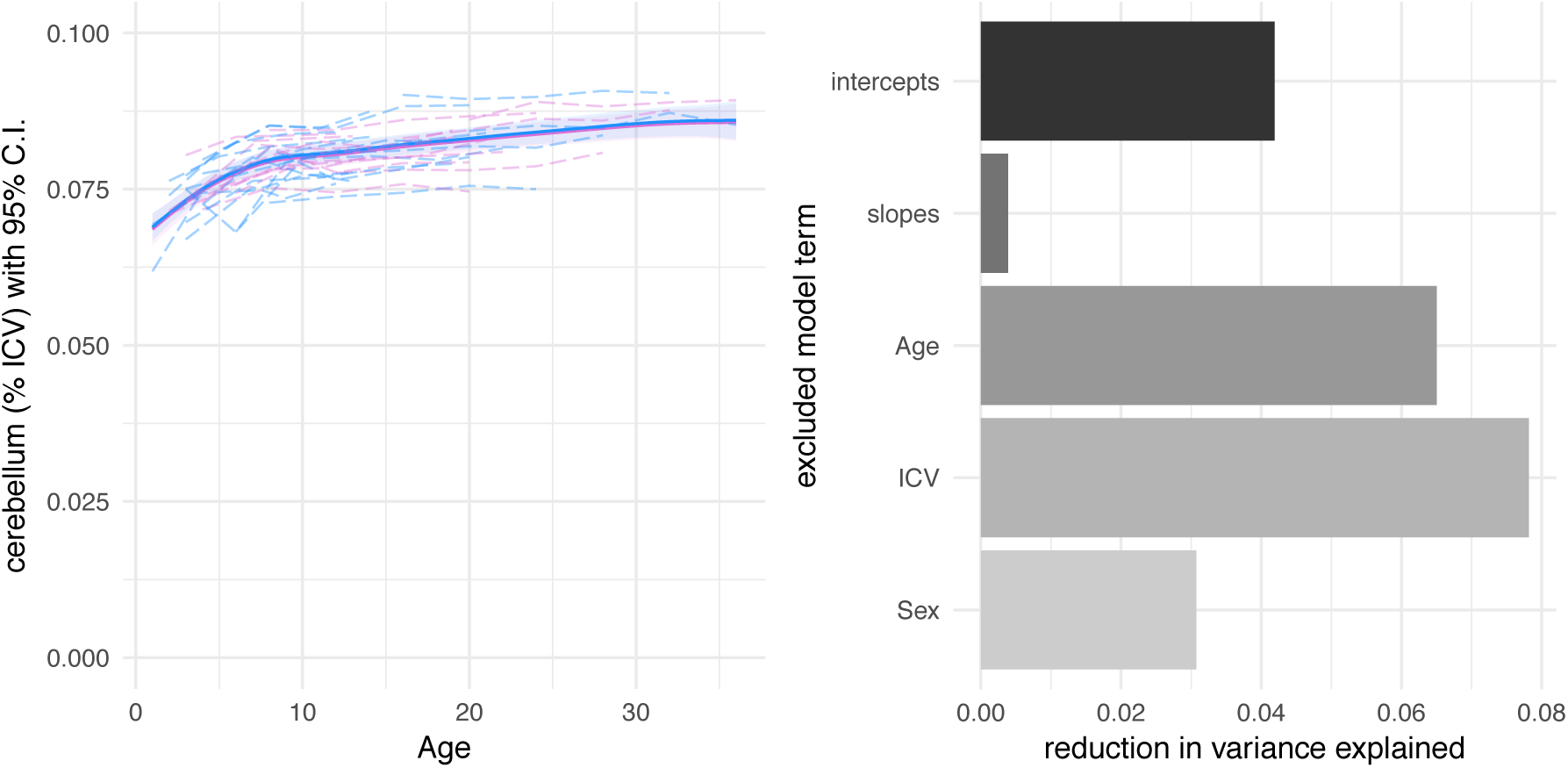
Longitudinal trajectories of cerebellar volume. Group average trajectories are visualised as a proportion of ICV for the cerebellum. Raw data are shown with dotted lines, solid lines represent model fit with 95% C.I. shaded. The reduction in variance explained when excluding model terms one-by-one from the full model fit is shown on the right. Trajectories for absolute volume are shown in Fig S2.

